# Quantifying individual influence in leading-following behavior of Bechstein’s bats

**DOI:** 10.1101/843912

**Authors:** Pavlin Mavrodiev, Daniela Fleischmann, Gerald Kerth, Frank Schweitzer

**Affiliations:** Chair of Systems Design, ETH Zurich, Weinbergstrasse 56/58, CH-8092 Zurich, Switzerland; Applied Zoology and Nature Conservation, University of Greifswald, Loitzer Strasse 26, 17489 Greifswald, Germany

## Abstract

Leading-following behavior as a way of transferring information about the location of resources is widespread in different animal societies. However, it cannot always be observed directly. Here, we develop a general method to infer leading-following events from observational data if only the discrete appearance of individuals is recorded. Our method further allows to distinguish such events from local enhancement at the resource, such as swarming behavior in case of bats, which is another widespread way of transferring information among animals. To test our methodology, we analyze longitudinal data about the roosting behavior of Bechstein’s bats from two different colonies and different years. The detection of leading-following events allows us, in a second step, to construct social networks in which nodes represent individual bats and directed and weighted links the leading-following events. We analyze the topology of these networks on the level of the colony, to see whether all individuals participate in leading-following behavior. Further, based on the leading-following network we measure the importance of individuals in this leading-following behavior by calculating different centrality measures. We find that individuals can be consistently ranked regarding their influence on others. Moreover, we identify a small set of individuals that play a central role in leading other bats to roosts. Our methodology can be used to understand the leading-following behavior and the individual impact of group members on the spread of information in animal groups in general.

## 1 Introduction

Leading-following behavior is prominent in different species to transfer information from informed to naïve individuals (Franks *et al*., 2002; Reebs, 2000; Kerth and Reckardt, 2003; Biro *et al*., 2006; Strandburg-Peshkin *et al*., 2015). Individuals who actively explore their environment, gather private information about the availability or the location of a certain resource, and subsequently lead naïve individuals to these resources (Franklin and Franks, 2012). By following a leader, naïve individuals gather information socially and become informed without having to spend prior search effort (Giraldeau and Caraco, 2000). When grouping at the resource is beneficial, e.g. during communal roosting, informed individuals benefit from leading naïve individuals as this increases the likelihood of conspecifics being present at the resource (Richner and Heeb, 1996).

This points to the question how individuals can assume their role as leaders or followers. Studies in collective motion have already reported that distinct leadership roles can emerge if some individuals are more active or better informed than others (Reebs, 2000; Pettit *et al*., 2013) or stand to gain more from imposing their preferences (Conradt and List, 2009; Rands *et al*., 2003). The presence of a small fraction of informed leaders has also been shown to be sufficient in guiding the movement of large groups with great accuracy in both human and animal societies (Couzin *et al*., 2005; Dyer *et al*., 2008). Some animal studies have even suggested that in addition to immediate cost and benefits, leadership is a personality trait independent of differences in information or knowledge of the environment (see Johnstone and Manica (2011) and references therein).

However, answering such primary questions becomes complicated when observations do not continuously track the information transfer through an animal system, but rather contain isolated individual measurements, e.g. discrete records of animal occurrences at measurement sites. In such cases, any leading-following behavior must be first reconstructed from the available data, for which one needs a sound methodology. It is one of the aims of this paper to provide this methodology, to (i) infer leading-following events from observational data and (ii) to distinguish such events from local enhancement at the resource. An example for such local enhancement is swarming behavior at potential day roost in the morning before bats collectively choose where to roost communally.

The second aim is to identify those individuals that play an important role in such leading-following behavior, by recruiting many other naïve individuals. This has as a precondition the reliable reconstruction of leading-following events from data. But it further needs an appropriate representation of the consecutive interactions between individuals and in particular a suitable measure to quantify importance, i.e. the influence on naïve individuals in leading-following behavior.

To reach this second aim, we build on the established methodology of social network analysis (Wasserman and Faust, 1994). Social network theory has transcended the human domain and has become widely accepted as an important conceptual framework for studying social interactions in animal groups (Croft *et al*., 2008; Wey *et al*., 2008; Pinter-Wollman *et al*., 2013). Its level of abstraction, where individuals become nodes and their interactions become links, allows us to quantitatively analyze social organisation in animal groups at all levels (individual, group, community, population, etc.) across a wide range of interaction types (recruitment, friendship, conflict, communication, etc.) (Krause *et al*., 2009).

As social structures in vertebrate animal systems are founded on behavioural interactions among individuals (Whitehead, 2008), social network analysis can be applied for studying social organisation in these systems as well. In this paper, we focus on Bechstein’s bats (*Myotis bechsteinii*), a forest-living, European bat species. During summer females form colonies that switch between many different communal day roosts in tree cavities and bat boxes (Kerth and Reckardt, 2003; Kerth *et al*., 2006; Fleischmann *et al*., 2013). In Bechstein’s bats, social network theory has unveiled the presence of long-term social relationships despite the high fission-fusion dynamics of the colonies, thereby imparting novel insights on the relation between cognitive abilities and social complexity (Kerth *et al*., 2011).

Specifically, in this paper we analyze the leading-following behavior of these bats to potential day roosts (bat boxes). After inferring such leading-following events from observational data, we construct a social network in which individuals are represented as *nodes* and their leading-following events as *directed and weighted links*, where the weights indicate the frequency of such events. This abstraction allows us to further analyze topological characteristics of such networks. On the level of the animal group (here, bat colony), this includes features such as connectedness, i.e. whether all individuals are part of the network. On the individual level, it allows to calculate centralities to infer the importance of the nodes, which translates to the influence of specific bats in this leading-following behavior.

To demonstrate the applicability of our methods, we analyze data sets from two different colonies of Bechstein’s bats and from five different years. This has implications for a better understanding of the collective behavior and information transfer about novel roosts in Bechstein’s bats. As we point out in the concluding discussions, we see the potential for a much broader application of our methodology to the leading-following behavior in different species.

## 2 Study animals: Bechstein’s bats

### 2.1 Coordination in roosting behavior

During summer adult female Bechstein’s bats form colonies to communally raise their young (adult males are solitary; Kerth and König (1999)). Such maternity colonies comprise 10-50 individuals, have a very stable individual composition, and are highly heterogeneous with respect to the age, reproductive status and the degree of relatedness among colony members (Kerth *et al*., 2002, 2011). Colonies switch communal roosts (tree cavities and bat boxes) almost daily and regularly split into several subgroups that use separate day roost (Kerth and König, 1999; Kerth *et al*., 2011). Communal roosting provides the females and their offsprings with grouping benefits, such as energetic advantages through clustering (e.g. social thermoregulation; Pretzlaff, Kerth and Dausmann, 2010; Kuepper, Melber and Kerth, 2016).

At the same time the frequent roost switching forces the female Bechstein’s bats to regularly explore new potential roosts during their nightly foraging trips and to coordinate their movements among day roosts in order to avoid permanent fission of the colony (Kerth and Reckardt, 2003; Kerth *et al*., 2006; Fleischmann *et al*., 2013). Experienced individuals, who have discovered the locations of suitable roosts through independent exploration, transfer their private knowledge to naïve conspecifics by leading them to these locations. Such leading-following events take place when one or several experienced bats arrive together with one or several naïve bats at a box at night. Information transfer about suitable roosts provides benefits to both the leading and the following bat. By leading conspecifics to potential roosts, an experienced individual increases the likelihood of communally roosting with conspecifics. At the same time, by following experienced individuals, naïve bats gather information socially without the need to spend prior search effort.

### 2.2 Field data collection

From 2007 to 2011, we studied two colonies (BS and GB2) of Bechstein’s bats within their home ranges located in two forests near Würzburg, Germany (Figure S1, left). Since 1996, all adult female bats in both colonies have been individually marked with individual RFID-tags in their first year of life (Kerth and van Schaik, 2012). Each RFID-tag is programmed with a unique 10-digit ID that can be identified and recorded by automatic reading devices Kerth and Reckardt (2003). The study period in each year was between the beginning of May and end of September. In that time, the colonies’ home ranges were equipped with about 20-30 experimental bat boxes per year in addition to a large number of already existing boxes (about 100; Fleischmann *et al*. (2013); Figure S1, right). These boxes were to serve as day roosts, similar to natural roosts in tree cavities, in which the Bechstein’s bats spend the day. All experimental boxes were equipped with RFID-loggers that recorded the bats’ nightly visits (Kerth and Reckardt, 2003; Fleischmann *et al*., 2013). In this way, every time a bat passes the entrance of an experimental box, its unique ID would be read and stored by the reading device without disturbance to the individual.

At the beginning of the study period in each year, the experimental boxes were placed within the home ranges and thus their locations were unknown to the bats until the first colony members discover them through private information gathering. Importantly, not all experimental boxes were discovered by the colony in a given year. Moreover, not all discovered and visited experimental boxes were subsequently used as day roosts.

Our datasets, thus, consist of the yearly recordings of the reading devices from all experimental boxes for each of the two colonies in each of the five years. Each recording contains a timestamp and the unique 10-digit ID of the bat who activated the reading device. An example dataset is shown in Table S1 in the supplemental material. Table 1 shows a summary of the total number of readings and the number of installed, discovered and occupied experimental roosts, for each colony throughout the years.

**Table 1:**
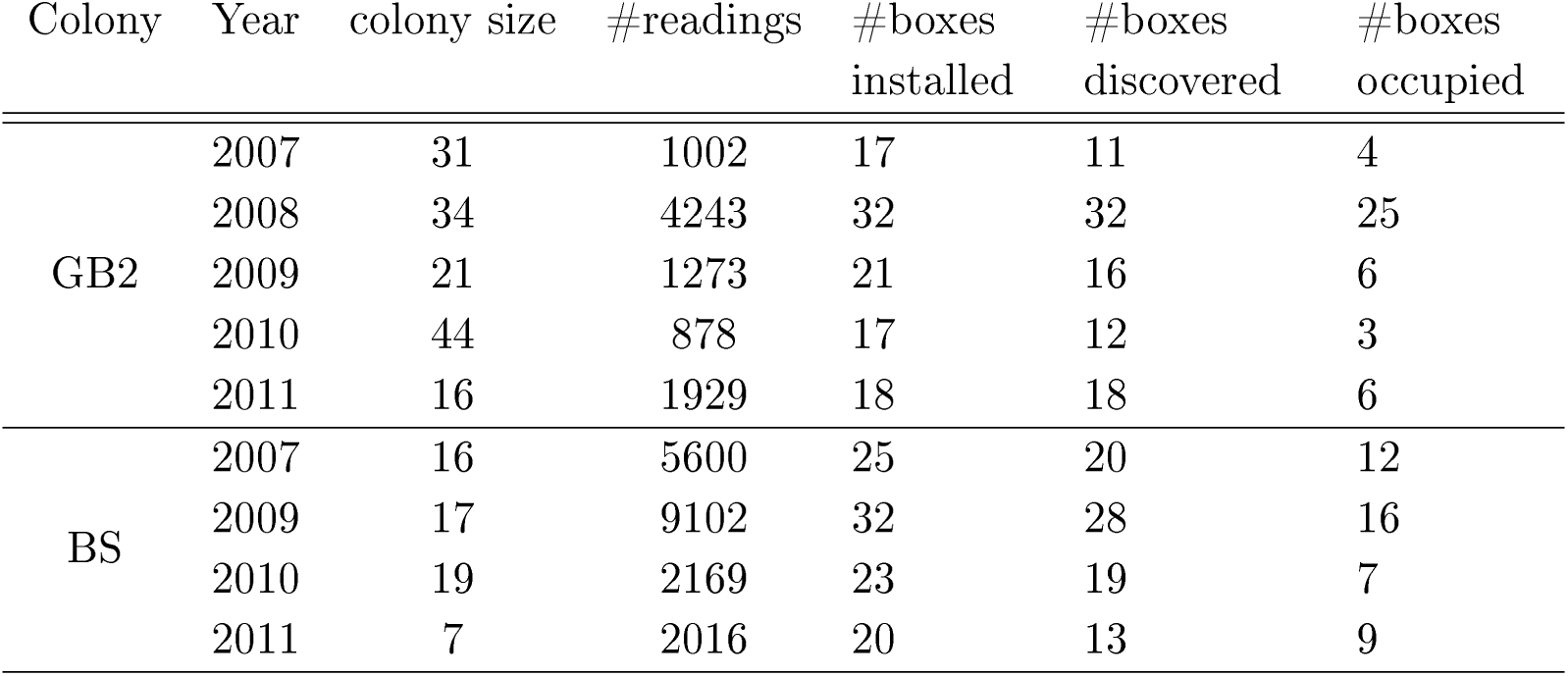
Data summary.

## 3 Methodology

### 3.1 Inferring leading-following networks

#### Defining leading-following events

Unlike studies on collective motion where group movement is tracked continuously (Strandburg-Peshkin *et al*., 2015; Ripperger *et al*., 2019), our datasets contain only discrete records of bat appearances at experimental boxes. Quantifying individual influence is, thus, contingent on a rigorous method for inferring leading-following events from discrete recordings of animal occurrences. To denote the information that individuals possess about the location of experimental boxes, we refine the nomenclature used by Kerth and Reckardt (2003). An individual bat is said to be naïve at time **t**_**1**_ regarding a given box, if it has *not* been recorded by the reading device in that box for all times **t** < **t**_**1**_. Similarly, an individual bat is considered experienced at time **t**_**2**_ regarding a given box, if it has been recorded in that box at any previous time **t** < **t**_**2**_. We define a *leading-following (L/F) event* to a given box at time **t**_**3**_ as the joint visit of two individuals - one naïve and one experienced at time **t**_**3**_. In case more than two bats arrive jointly, we form all possible L/F pairs consisting of one naïve follower and one experienced leader.

With this definition of L/F events, the actual inference of L/F event patterns from the data relies on three parameters: (1) the maximum allowed time difference (in minutes) between consecutive recordings of a leader and a follower, (2) the minimum time (in minutes) an experienced bat in an L/F event needs to potentially become a leader, i.e. the time needed to find and lead followers, and (3) the hour in the morning on the day of a box occupation, after which subsequent recordings from this box are ignored because of swarming behavior (local enhancement). In Sections S.3 and S.4 of the Supplementary Material we present the explanation for these parameters together with a rigorous statistical procedure for their calibration.

#### Constructing leading-following networks

Following the above procedure, we identified all L/F events in each of our datasets. We then constructed directed and weighted leading-following (L/F) networks, aggregated over the duration of the study period. In these networks, a node represents an individual bat and a link between two nodes indicates their involvement in a leading-following event. More specifically, links are *directed*. A directed link from node A to node B, denoted as A → B, means that individual A followed individual B to a given experimental box. The weight of this directed link is the number of times that A followed B (to different experimental boxes) during the study period.

We also compute the number of weakly connected (WCC) and strongly connected components (SCC). A WCC of a network is a sub-network in which any node can be reached from any other node, either by a link between these two nodes, or by following a sequence of links through other nodes, regardless of the direction of these links. Similarly, a SCC is a WCC with the additional restriction that the *direction of the links* must be respected when connecting any two nodes. As we explain in the next section, these two measures are particularly important for judging the extent to which information can spread in a network.

### 3.2 Social Network Analysis

#### Quantifying individual influence

Social network analysis builds on the existence of a social network that can be analyzed. Such a network has been constructed in the previous step, where directed links represent leading-following events between individual bats. We can now use the *topology* of the network, i.e. the relation between nodes expressed by their links, to characterize the position of individuals in such a network.

Our aim is to identify those nodes, i.e. individual bats, that are most influential in leading other bats. In social network analysis, the importance, or influence, of a node in a certain dynamical process flowing through the network is referred to as *centrality*. There are various centrality measures in use, and each makes certain implicit assumptions about the dynamical process flowing through the network (Borgatti, 2005). Choosing a centrality measure is, thus, context-dependent (see Figure 1).

**Figure 1:**
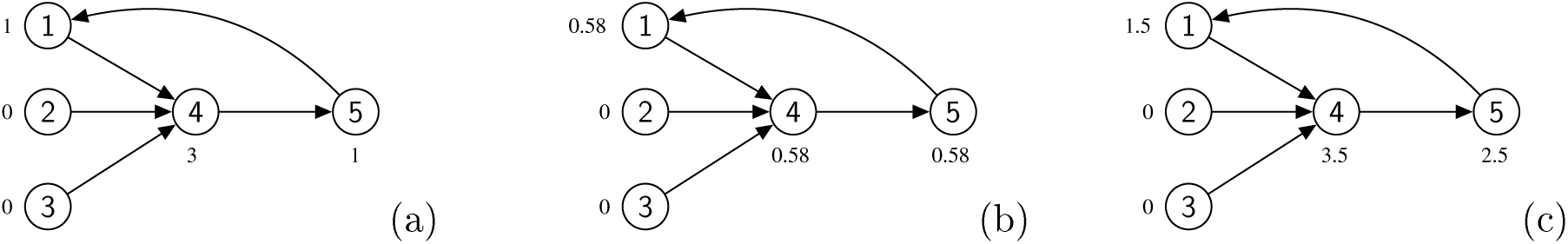
Differences between the three candidate centrality measures. The centralities for each measure are indicated next to each node. **(a)** *In-degree centrality*. Here, only direct influence is measured. Individual 4 is most influential, as she spread information to three different individuals. Individuals 1, and 5 with one follower each, have still equal importance. **(b)** *Eigenvector centrality*. Since individuals 2 and 3 have no followers, they are attributed zero influence, and thus contribute nothing to the influence of their leader, individual 4. In turn, 1, 4, and 5, each have one follower of non-zero importance, hence they have the same eigenvector scores. **(c)** *Second-degree centrality* with *α* = 0.5. Individual 4 has a higher centrality than her in-degree score, as we account for the indirect contribution of individual 1 (3 + 0.5 × 1 = 3.5). However, 5 is now more important than 1, because 4 contributes to 5 indirectly (1 + 0.5 × 3 = 2.5).

#### In-degree, eigenvector and second-degree centrality

In our case, an appropriate centrality measure must reflect the notion of individual importance in spreading information about suitable roosts. The simplest possible measure is the *in-degree* centrality (Figure 1a), which defines individual importance as the total number of bats that an experienced bat spread information to directly. In-degree centrality is, thus, calculated as the weighted sum of all directed links that point to a given experienced individual.

In-degree centrality measures the total number of leadings, i.e. direct influence, *without* considering how the information distributed by a leader to its followers propagates further through the colony. To also account for such indirect effects, an alternative centrality measure is *eigenvector centrality* (Figure 1b). In a social network, a node has high eigenvector centrality if it is pointed to by nodes that themselves have high eigenvector centralities. In other words, an experienced bat leading a few bats, who themselves lead a lot can be more influential than a bat leading many other bats who never lead. The computation of eigenvector centralities is presented in Section S.5 of the Supplementary Material.

The in-degree and eigenvector centralities represent two extremes, the former measuring exclusively direct influence, and the latter additionally measuring all possible indirect ways, in which information can flow from one individual to all the rest. Eigenvector centrality, however, considers all chains to be of equal importance. Hence, this metric will grow with the length of the chain and individuals who are part of longer chains will tend to be quantified as more influential. This influence, however, does not reflect genuine information spreading, as it is quite likely that beyond length two, the target roost of the L/F events further down the chain, changes.

To address this issue with eigenvector centrality, we define a *new metric - second-degree centrality* (Figure 1c) - which computes centrality as the in-degree of the focal individual and the sum of the in-degrees of its followers, weighted by a factor *α* (in that sense the followers of one’s followers are its second-degree followers). This reflects our observation that chains of length up to two constitute the majority in all datasets. We, thus, use second-degree centrality as the main measure for quantifying individual influence.

## 4 Results

### 4.1 Chains of L/F events

Using the above definition of L/F events, first we have determined the three relevant parameters to determine L/F events in the data as (1) 5 minutes for the maximum allowed time difference, (2) 3 minutes for the minimum time, and (3) 5am the hour in the morning on the day of a box occupation (see Sections S.3 and S.4 of the Supplementary Material for a detailed explanation and a statistical calibration).

Identifying all L/F events allows us to construct the respective network in the following. Before, however, we are interested in the occurence of *chains* of L/F events of a certain length, through which information about a fixed roost is spread. For example, two L/F events, A → B and C → A, constitute a chain of length two (in addition to forming two separate chains of length one), provided both were to the same roost. In other words, we assume that B spread the information to A, and A, in turn, transferred it further to C. Therefore, B ought to obtain direct importance from having led A, but also indirect contribution, for were it not to B, A would not have learned about this box and thus could not lead C to it. This assumption is not entirely correct, however, since it is possible, though unknowable, that A would have found the roost by its own exploration, or that A “forgot” the information obtained from B, and re-visited the box before leading C. The latter issue is exacerbated with the length of the event chains we consider.

Figure 2 shows the relative frequency, aggregated over all datasets, of observing chains of L/F events. This frequency can be interpreted as the probability of finding chains of a given length. As the inset in Figure 2 demonstrates, the probability distribution resembles an exponential distribution. The plot further indicates that chains longer than 16 did not occur in any of the datasets we have. More importantly, event chains of length up to two constitute about 80% of all lengths observed, and the probability of longer chains decreases drastically. We, thus, argue that the long L/F chains we observe in the L/F networks likely do not represent information spread about the same roost, and should therefore be discounted by any influence measure.

**Figure 2:**
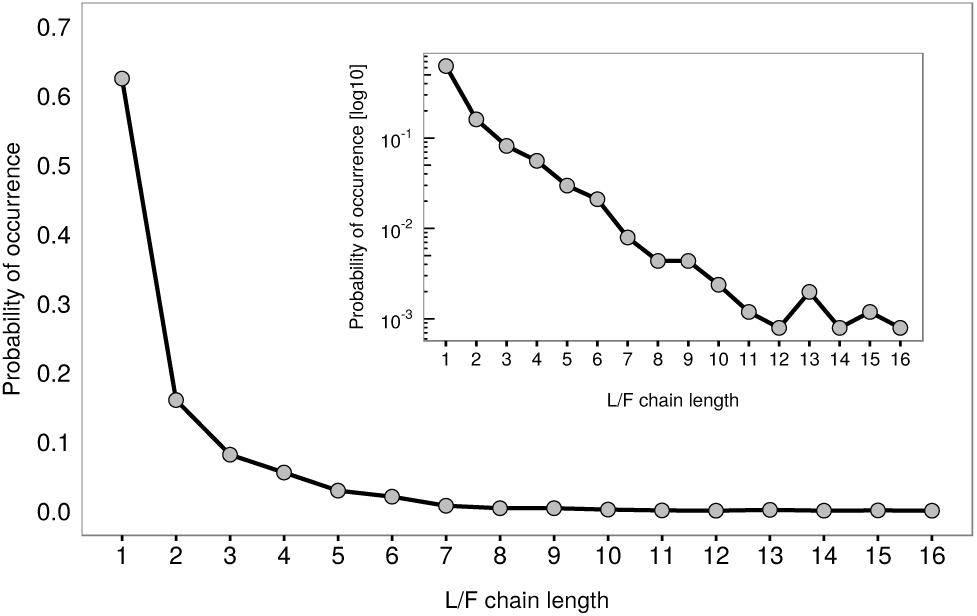
Probability distribution of the lengths of L/F event chains, calculated over all nine datasets. Inset: log-linear plot of the data

### 4.2 Constructing the L/F network

As explained, the L/F network illustrates all detected L/F events, where nodes represent individual bats and directed links represent leading-following events. The data is aggregated over time, thus the width of the links indicates the number of events in the dataset. In Figure 3 we illustrate the L/F network for the GB2 colony in the year 2008. The reasons for concentrating on this colony are discussed in Section S.3. Looking at Figure 3, we realize that individuals differ remarkably with respect to their importance, as reflected both by their in-degree centrality (size of the nodes) and their eigenvector centrality (node color). It is also evident that there are correlation between in-degree and eigenvector centrality, as visible for the four individuals in the center.

**Figure 3:**
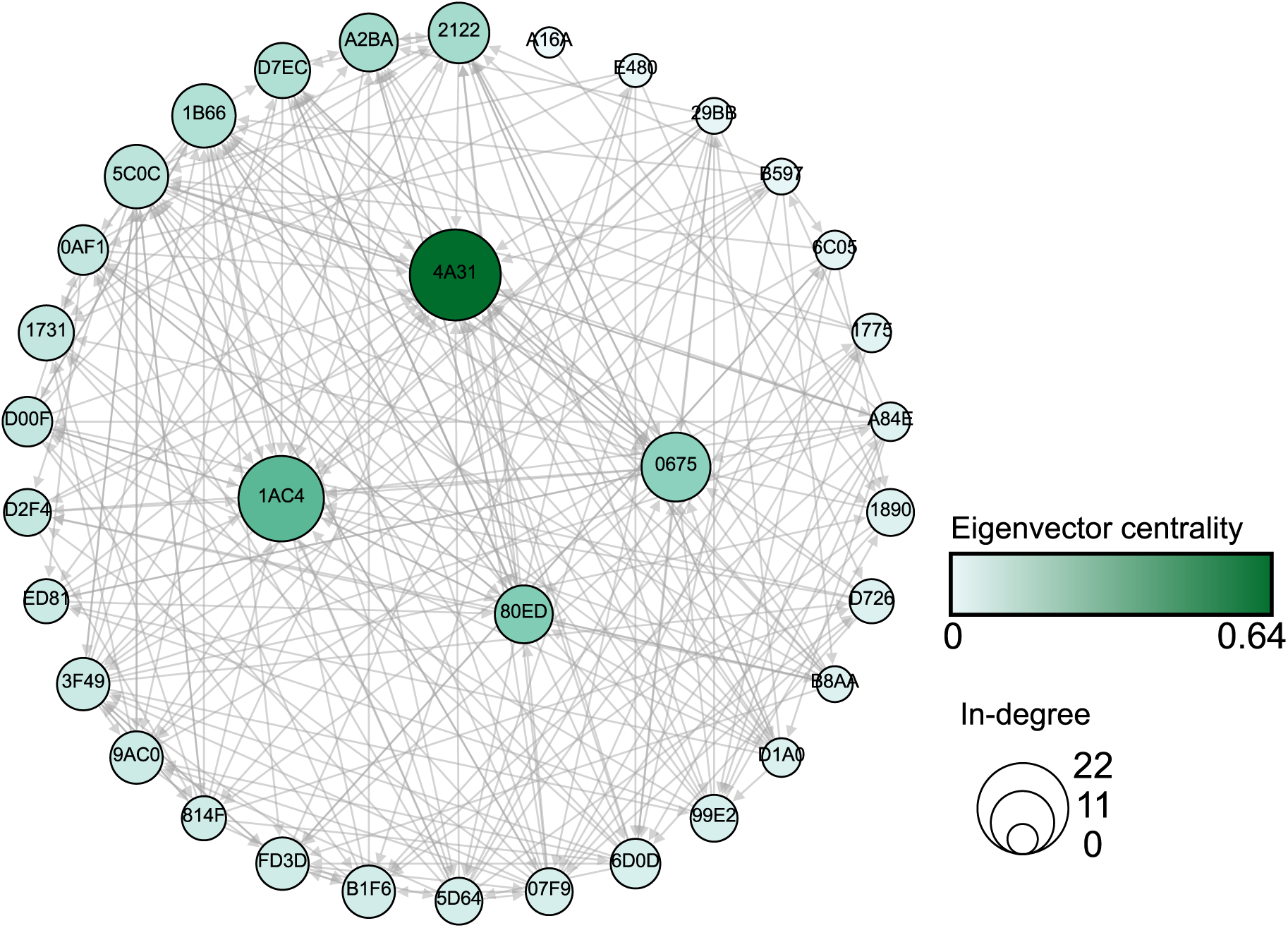
Aggregated leading-following network for the GB2 colony in 2008. Nodes represent individual bats (indicated by a hexadecimal number inside the circle). Directed links represent following behaviour. Node colors indicate eigenvector centrality, whereas node sizes indicate in-degree centrality. The four individuals with highest eigenvector centrality are shown in the middle. Note that for the sake of illustration, links show only *unique* L/F events. I.e. leading-following between the same leader and follower, but to different roosts, are omitted to maintain the readability of the graph. Total number of unique L/F events is 262, while the total number of L/F events, including multiple leading-following between the same individuals, is 321 (Table 2).

Table 2 presents salient network characteristics, regarding the degree of connectedness of the L/F networks *in all datasets*. Network density is defined as the fraction of inferred L/F events out of the maximum possible number of L/F events for that network. For example, the L/F network for the GB2 colony in 2007 consists of 31 individuals, hence the maximum possible number of L/F events is 31 × 30 = 930, which yields a network density of 0.06.

**Table 2:** Topological characteristics of the leading-following networks from the GB2 and BS colonies. Shown are number of bats (nodes), number of identified L/F events (links), network density, number of weakly connected components, number of strongly connected components, and the size of the largest strongly connected component. Rows in cyan are the dataset we consider for further analysis.

| Colony | Year | #bats | #L/F events | density | #WCC | #SCC | size of largest SCC |
| --- | --- | --- | --- | --- | --- | --- | --- |
| GB2 | 2007 | 31 | 60 | 0.07 | 4 | 23 | 9 |
|  | 2008 | 34 | 262 | 0.23 | 1 | 2 | 33 |
|  | 2009 | 21 | 33 | 0.08 | 2 | 19 | 3 |
|  | 2010 | 44 | 142 | 0.08 | 1 | 22 | 14 |
|  | 2011 | 16 | 86 | 0.35 | 1 | 2 | 15 |
| BS | 2007 | 16 | 169 | 0.70 | 1 | 1 | 16 |
|  | 2009 | 17 | 201 | 0.74 | 1 | 1 | 17 |
|  | 2010 | 19 | 148 | 0.43 | 1 | 3 | 17 |
|  | 2011 | 7 | 26 | 0.62 | 1 | 1 | 7 |

We see that the two colonies differ in this respect through the years. While the L/F networks for the BS colony displays high density and connectivity for all study years, the L/F networks for GB2 colony in the years 2007, 2009 and 2010 have low density consistent with the fewer L/F events observed. Therefore, to calculate the importance of each individual, we use only the cyan-coloured datasets in Table 2, as they provide the most reliable sample sizes of detected L/F events for statistical analysis.

If we focus only on these datasets, we find that their respective L/F networks are weakly connected. I.e. there is only *one* weakly connected component, which means that all individuals participated in L/F events. Moreover, these networks consist of only a few (1-3) strongly connected components (SCC). Within an SCC, each individual can be reached from any other individual by following (a chain of) *directed* links. In most of the chosen cases, the size of the largest SCC is similar to the total number of nodes, which means that the vast majority of individuals participated as *both* leaders and followers. Otherwise, one could reach a given individual through a directed chain, but will not be able to connect from this individual back to the network via a directed chain. Hence, individuals would be part of a weakly connected component (WCC) because they are *either* followers *or* leaders, but they would not be part of a SCC.

### 4.3 Quantifying individual influence

The construction of the different L/F networks as described above now allows us to quantify the importance of individuals in these networks. For this, we use the three different centrality measures introduced in Section 3.1, i.e. in-degree centrality, eigenvector centrality and second-degree centrality.

Figure **S2** in the Supplementary Material shows the results of each of these measures separately for the colony GB2 for the year 2008. If we compare the *absolute values* of the centralities, we find that influence scores are heterogeneous with a *majority* of individuals exerting low to mid influence and a *minority* having high influence. This result holds regardless of the centrality metric used to quantify influence. We note that already the visualization of the L/F network in Figure 3 uses the information of centrality values.

We can use the absolute values to determine the *relative* importance, by ranking individuals according to their second-degree centrality. The results are shown in Figure 4, where the diagonal indicates increasing rank, i.e. decreasing importance. In order to determine whether these results are robust if instead of second-degree centrality the other two measures are used for the ranking, we have provided the respective ranks in the same plot. As Figure 4 shows, the three proposed centrality measures produce a highly consistent ranking of individual influence.

**Figure 4:**
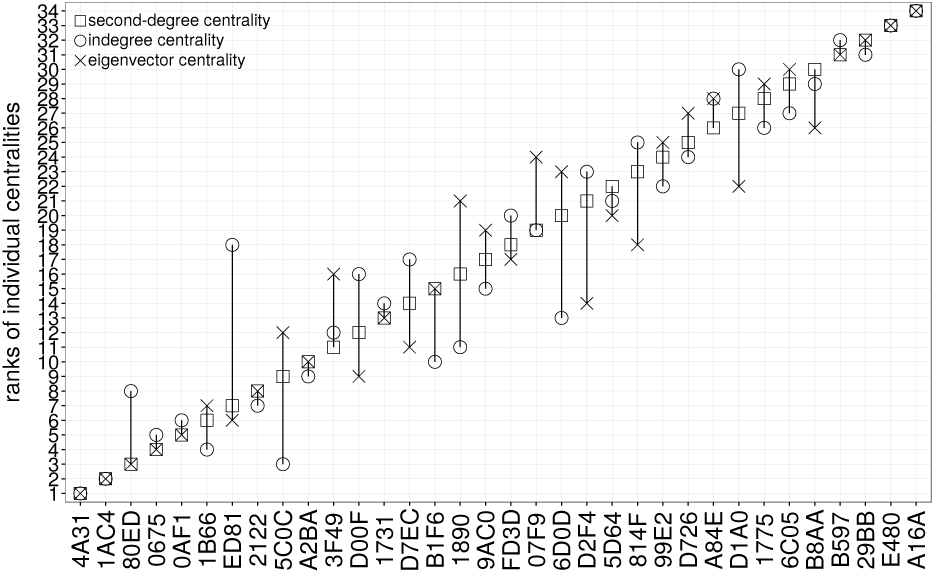
Ranked individual influence of bats of colony GB2 in year 2008. The x-axis displays the last four digits of a bat’s unique identification number. The y-axis displays the rank according to the second degree centrality (square symbols) in increasing rank order (rank 1 - highest centrality). For each bat we additionally plot its rank when importance is quantified as in-degree (circle symbols) and eigenvector centrality (cross symbols). Overlap of the three symbols indicates that the given individual has the same rank, regardless of the centrality measure used. For the individual centrality values see Figure **S2**.

To verify this finding, we have extended the above analysis to all datasets indicated in Table 2. For each dataset, we have then calculated the Pearson correlation between the rankings obtained from the three centrality measures. The results are given in Table **S10** in the Supplementary Material. We find that for all datasets the Pearson correlation is very high for all combinations. That means that ranking individuals according to any of the measures leads to a consistent rank of influence.

## 5 Discussion

This paper provides a general methodology for inferring interaction networks from proximal data. We use rich longitudinal data sets of joint visits of Bechstein’s bats in potential day roosts. While proximal networks do not always correlate well with interaction networks (Castles *et al*., 2014), it has been argued (Farine, 2015) that proximity is a good proxy for interactions in fission-fusion societies such as Bechstein’s bats.

Below we summarize our approach as it differs from common techniques (Farine and Whitehead, 2015) to study animal association patterns via social networks. Typically, when social networks are used, the *observed interaction strength* between two individuals is either thresholded, sampled or used as a link weight, to calculate various association indexes (Franks *et al*., 2010; Croft *et al*., 2008; Hoppitt and Farine, 2018). In line with Farine and Whitehead (2015), we do *not* threshold our networks to avoid dubious statistical biases. Instead, we include *all* of the observed individuals and their recorded activity and analyze the *full scale* of inferred interactions.

Moreover, we also go beyond calculating association indexes and the corresponding Mantel tests. Association indexes are *local* measures in that they only reflect dyadic relations between any two individuals. To quantify the *systemic* influence of individuals, we need to provide measures that also capture their proclivity to act as social hubs, as recognized already by Farine and Whitehead (2015) and Brent (2015). Therefore, we have proposed a novel *centrality measure*, second-degree centrality.

Our methodology builds on raw data that contains only the recordings of single bats entering a given roost site at a particular time. Such data *per se* does not contain any information about importance, or influence. We focus on a specific type of influence, namely that an experienced individual leads an inexperienced, i.e. naïve, individual to a particular roost. Therefore, the first challenge is to identify leading-following (L/F) *events* from this data and to construct a social *network* from all these L/F events, and the second challenge is to quantify the *importance* of individuals in this leading-following network, appropriately.

Regarding the first challenge, we note that most field experiments, including ours, are limited by the state-of-the-art passive RFID-tagging, which only records presence data. There is a more advanced technique (Ripperger *et al*., 2019) that uses an proximity sensor system to continuously track the leading-following behaviour between female bats and their juvenile to suitable roosts. However, such technology is still in its nascent stage and not widely used in field experiments, as with this battery-powered system small bat species cannot be tagged at present and it is not possible to follow many individuals over an extended period of time.

Our methodological contribution can be also adopted for other species where leading-following behavior plays a role and only recordings of individual positions are available. This includes, for example, automatic RFID-tag recordings at feeding stations (Farine and Whitehead, 2015) and other resources where different group members meet, such as burrows in rodents (König *et al*., 2015). As we demonstrate, such recordings can be systematically analyzed by comparing (statistically) the distributions of L/F time differences, to infer genuine L/F events.

A major contribution of our analysis is a thorough investigation of the parameters that allow to distinguish a L/F event from other types of encounters (e.g. local enhancement) at a given box. We recall that there is no ground truth available that tells us about the correct identification of L/F events from the data. We argued that the time differences of L/F events can be used to calibrate three relevant parameters: (1) the maximum allowed time difference (in minutes) between consecutive recordings of a leader and a follower, (2) the minimum time (in minutes) an experienced bat in an L/F event needs to potentially become a leader, i.e. the time needed to find and lead followers, and (3) the hour in the morning on the day of a box occupation, after which subsequent recordings from this box are ignored because occupation is considered to have already taken place. For a rigorous statistical analysis of the influence of these three parameters on inferring L/F events see the Supplementary Material.

Regarding the second challenge, we have proposed a new measure of individual influence that can be derived from these L/F events. Obviously, there is no natural distinction between only leaders and only followers in the observed bat colonies. Instead, almost all individuals are *both* leaders and followers, but at different times and, importantly, to a different degree. For comparison, in African elephants, a single matriach leads a group and the group members profit from following her as she has long-term experience (McComb *et al*., 2001). In primates, the individual influence of group members can depend on the context, and may range from a single dominant individual who influences where a group moves to, to a more widely distributed influence on travel destinations among group member (King *et al*., 2008; Stueckle and Zinner, 2008).

To quantify individual influence, we have constructed a *social network* in which nodes represent individual bats, directed links indicate a leading-following event and the weight of the links considers the frequency of such events. Analyzing the topology of these social networks already allows us to draw several conclusions about the information sharing in the respective colonies.

First, note that we focus our analysis on dense networks (see Table 2). We found that these networks have only one weakly connected component (WCC) which contains most of the individuals. Density is a proxy for the intensity of leading-following behaviour, while the presence of one large WCC indicates that the majority of the colony partook in leading-following. Moreover, we also found that in most cases there are only very few (1-3) strongly connected components (SCC) of different size in the network (see Table 2). Hence, we can conclude that individuals in the same SCC participated both as leaders and as followers in different events. This tells us that information about suitable roosts is not concentrated in only a few important individuals, but is spread across the whole colony.

At the same time, we could also detect that not all individuals play an *equal* role as leaders or followers. Instead, their influence, measured by leading inexperienced bats, differs considerably. To quantify these differences, we used different centrality measures as proxies of importance. Two of these, in-degree and eigenvector centrality, are established measures, while the third one, second-degree centrality is a new measure introduced by us. As explained in Section 3.2, it cures certain shortcomings of the other two centrality measures if applied to L/F networks. When considering aggregated measures, such as *rankings*, second-degree centrality is correlated to in-degree and eigenvector centrality, because it is derived from them. However, second-degree centrality differs on the individual level, as it more accurately reflects the genuine information spreading observed in the data.

Computing the different centralities for each individual, we could identify that there are only a few important individuals that lead most of the other bats. These individuals stand out regardless of the centrality measure used. In particular, we also calculated that there are significant correlations between the rankings obtained by using the different centrality measures. We emphasize that measuring influence by means of centralities cannot be simply reduced to comparing numbers of leading events. The latter would not allow us to distinguish whether individuals always lead the same or diverse followers, or whether such followers are of less or equal importance in comparison to the leader.

We believe that our results can guide future empirical and theoretical studies in two ways. First of all, we should realize that the constructed L/F networks do not already tell us about the *mechanisms* by which pairs of leaders and followers are formed. This process, known as *recruitment*, can be revealed by testing different recruitment rules in computer simulations, to check whether they result in the importance scores obtained from the empirical networks. In essence this entails the development of various *null models*. Null models are recognized as useful tools to test the viability of these recruitment rules in the presence of inherently non-independent behavioral data (Farine, 2017). We investigate a variety of such null models about recruitment behavior in Bechstein’s bats in a subsequent paper (Mavrodiev *et al*., 2019).

Secondly, additional field work needs to be devoted to study the behavioural variability of individuals in playing their role as leaders or followers. For example, demographic, health or genetic characteristics can influence such roles (Brent *et al*., 2015; Fischhoff *et al*., 2007; Keiser *et al*., 2016; McComb *et al*., 2011). With our study, however, we have already identified those individual bats that are prominent in these roles. This allows to target future experiments particularly toward individuals with very high or very low influence, to find out how different characteristics impact their leading-following behavior.

## Acknowledgements

We thank the local forestry and conservation departments for their continuous support throughout this long-term study, and the numerous people who helped gather the field data used in this study, in particular Anja Baigger and Markus Melber. This work profited strongly from the financial support of the German (DFG, KE 746/2-1/3-1/4-1/5-1/6-1) and the Swiss National Science Foundation (SNSF, 31-59556.99) during the years covered in this study.

## Electronic Supplementary Information

### S.1 Illustration of the raw recordings in our datasets

**Table S1:**
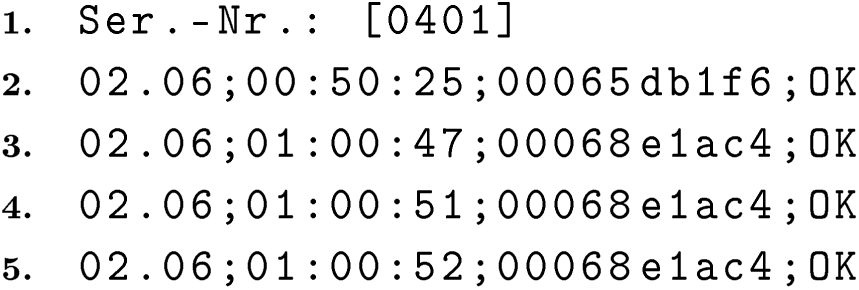
An excerpt from the recordings of an experimental box for the GB2 colony in year 2008. Line numbers serve as a visual guide only and are not part of the data. Each line corresponds to one reading, i.e. one activation of the reading device by a visiting bat. Columns are separated by semicolon. The first column shows the date of the reading (in this case June 2^nd^), the second column indicates the time of the recording in 24-hour format, the third column contains the unique 10-digit ID of each bat, and the last columns is a status message.

### S.2 Inferring L/F events

Recall that an L/F event is defined as the joint visit of an experienced and a naïve individual at a given box. Furthermore, we associate with each L/F event the experimental box in which it was detected, and the times at which the leader and the follower were recorded by the reading device in the box. Note that, it is not necessary for the leader to enter the box before the follower. Often it is the latter who is registered first. In case the leader and the follower were recorded multiple times, we take those times that minimize the difference between their appearances in the dataset (see Table S2 and associated explanation). Finally, we refer to the **time_difference** of an L/F event as the absolute difference between the recording times of the leader and the follower.

The actual inference of L/F events from the definition above depends on *three parameters*. The *first* parameter is the *maximum time difference* allowed between consecutive recordings of a leader and a follower, regardless of order. We refer to it as **lf_delay**. The **lf_delay** is important in determining which patterns constitute a joint visit of two individuals, as bats do not enter a box immediately upon arriving: females returning at night to a day roost usually encircle it several times before entering (Kerth and Reckardt, 2003; Schöner *et al*., 2010). Therefore, **lf_delay** limits the sheer number of L/F events we detect, since the higher the limit, the more likely it is to find an experienced and a naïve individual recorded within **lf_delay** of each other. In the limit of **lf_delay** → *∞*, we would detect the maximum number of L/F events, many of which would be false positives, as bats recorded days apart would still be assumed to have “jointly” arrived at a box.

The *second* parameter represents the *minimum* time a follower in an L/F event needs to potentially become a leader, i.e. the time needed to find, recruit, and lead other followers. We denote it as **turnaround_time**. The importance of this parameter becomes apparent in Table S2, which shows a frequently occurring recording pattern.

**Table S2:**
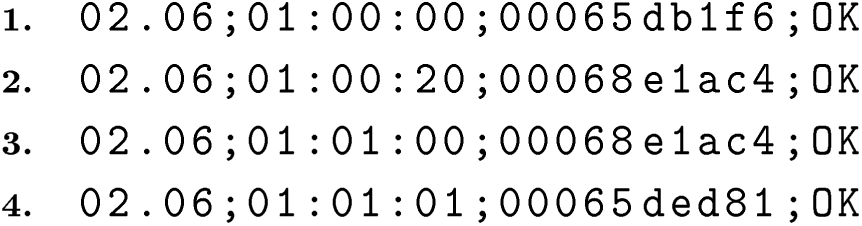
A simplified example of how turnaround_time affects the inference of L/F events.

Assume that, for this box, individual **00065db1f6** is experienced at time **01:00:00** (line 1), individual **00068e1ac4** is naïve at **01:00:20** (line 2), and individual **00065ded81** is naïve at **01:01:10** (line 4). Taking **lf_delay**=**3** minutes (which is a good rule-of-thumb Kerth and Reckardt (2003) we can deduce that individual **00068e1ac4** followed individual **00065db1f6** to that box, i.e. **00068e1ac4**→**00065db1f6**. More precisely, we infer an L/F event to this box with the leader recorded at **01:00:00** and the follower at **01:00:20**. The time difference of this event is 20 seconds.

Let us further assume that **00068e1ac4** liked the box she was just led to, and in turn would like to show it to other individuals. Its second recording in this dataset is on the third line - 40 seconds after its first appearance as a follower. If we assume that **turnaround_time** < 40 seconds, then we also have to assume that **00068e1ac4** would have had enough time to fly within its home range, meet other individuals, recruit and ultimately lead them back to this box. In this example, she led individual **00065ded81** who appeared within a time of **lf_delay** from it, i.e. we then also have to infer the L/F event **00065ded81**→**00068e1ac4**. In addition, however, we see that **00065db1f6** and **00065ded81** appear within **lf_delay** of each other, hence we must also form the L/F pair **00065ded81**→**00065db1f6**. Evidently this last L/F event contradicts **00065ded81**→**00068e1ac4**. Hence, **turnaround_time** < 40 seconds is a wrong assumption.

The issue is that, in reality, the 40-second delay between the two readings of **00068e1ac4** is most likely not due to it having led another individual to the box. Instead, it is highly likely that either (i) the first reading showed the bat entering the box and then leaving it again shortly thereafter or (ii) that the bat was simply encircling the box for 40 seconds, and then triggered the reading device a second time upon re-entry. The proper distinction between actual recruitment and such behavioural variability is the role of the parameter **turnaround_time**. In the toy example from Table S2 a more realistic interpretation is that **00065db1f6** led both **00068e1ac4** and **00065ded81**, i.e. we would only infer two L/F events. Note that since **00068e1ac4** appears twice, we associate the time of its first recording (**01:00:20**) with the L/F event **00068e1ac4**→**00065db1f6**, since it minimizes the time difference to the recording of the leader.

The *third* parameter is the hour in the morning, on the day of a box occupation, after which subsequent recordings from this box are *ignored*. The necessity to ignore some recordings comes from the need to distinguish between genuine information exchange about suitable roosts (in terms of leading-following) and “pre-occupation” behaviour. Before the occupation of a given box, experienced individuals who have decided to roost there, fly around the box and emit echolocation calls that attract naïve individuals to the same box (O’Shea and Vaughan, 1977; Schöner *et al*., 2010). It has been suggested that this broadcasted information is used by naïve bats (especially juveniles) to learn the location of suitable roosts from experienced conspecifics (Kerth *et al*., 2003). The result is that occupation is preceded by a growing group of individuals (experienced and naïve) flying around, or *swarming*, the roost for several hours. In our data, this is reflected by readings of naïve individuals, which appear shortly after each other in a long sequence, together with the readings of experienced bats. As a result, additional L/F events will be identified with time differences close to the allowable limit of **lf_delay** (see Section S.4 for illustration). These L/F events do not constitute genuine recruitment, in the sense that naïve individuals were led to a roost, but rather reflect the swarming phenomenon (local enhancement). Therefore, we define the parameter **occupation_deadline** as the temporal deadline on the day of a box occupation, after which subsequent readings in this box are attributed to swarming, and thus ignored.

### S.3 Selecting parameter values

As illustrated in Section S.2, each of the three parameters affects the inference of L/F events differently. Therefore, it is important to choose proper values that allow us to identify an adequate number of genuine leading-following events for statistical analysis. Empirical research in the field of information transfer in Bechstein’s bats has suggested 3 minutes for **lf_delay** and 3am for **occupation_deadline** as a reasonable rule of thumb (Kerth and Reckardt, 2003). We build upon these heuristics by comparing the distributions of time differences of all L/F events, fixing **lf_delay** and varying the other two within a reasonable range (see Figure **S1**).

To generate sufficient sample sizes for the comparison, the dataset we chose to analyze was the GB2 colony in 2008 (Table 1 in main text). The reason is that, in 2008, the colony had the highest number of discovered and occupied boxes, the second largest colony size, and a large amount of individual readings. Therefore, we expected to identify the largest number of L/F events from this dataset. Note that any combination of the three parameters is a 3-tuple, which generates a set of L/F time differences from all identified L/F events in the dataset. An example is presented in Figure **S1**, where we show histograms of the L/F time differences for **lf_delay-turnaround_time**=3 minutes, and **occupation_deadline**=2am (left) and **occupation_deadline**=3am (right). Figure **S1** also illustrates why we focus on the distributions of L/F time differences to select the values of the three parameters. As there is no objective method ^†^ to quantify the behaviour underlying each of the parameters, we argue that L/F time differences best capture the effect that varying the parameters has on the L/F events we identify. For example, a visual inspection of Figure **S1** hints that increasing **occupation_deadline** from 2am to 3am does not change the time difference distributions. This implies that swarming has not yet set in (otherwise, we would expect quantitatively more events with longer time difference), and the additional L/F events on the right-hand side are genuine. Consequently, we would prefer **occupation_deadline**=3am, as it increases our sample size. Table **S3** formalizes this argument.

**Figure S1:**
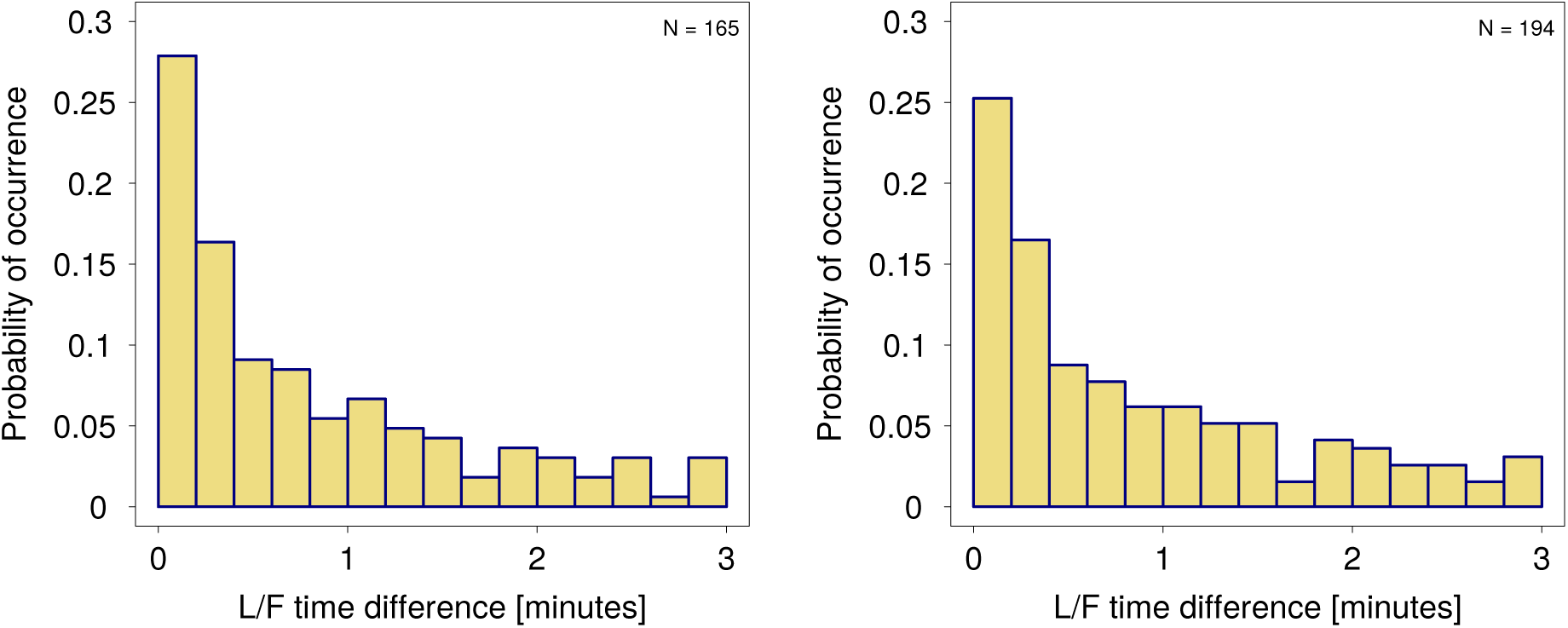
L/F time differences for the GB2 colony in 2008. Histograms show the absolute differences between the times at which the leader and the follower were recorded in all identified L/F events. Parameters: `turnaround_time` = `lf_delay` = 3 minutes (both plots), `occupation_deadline` = 2am (left) and `occupation_deadline` = 3am (right). Insets indicates the total number of identified L/F events.

**Table S3:**
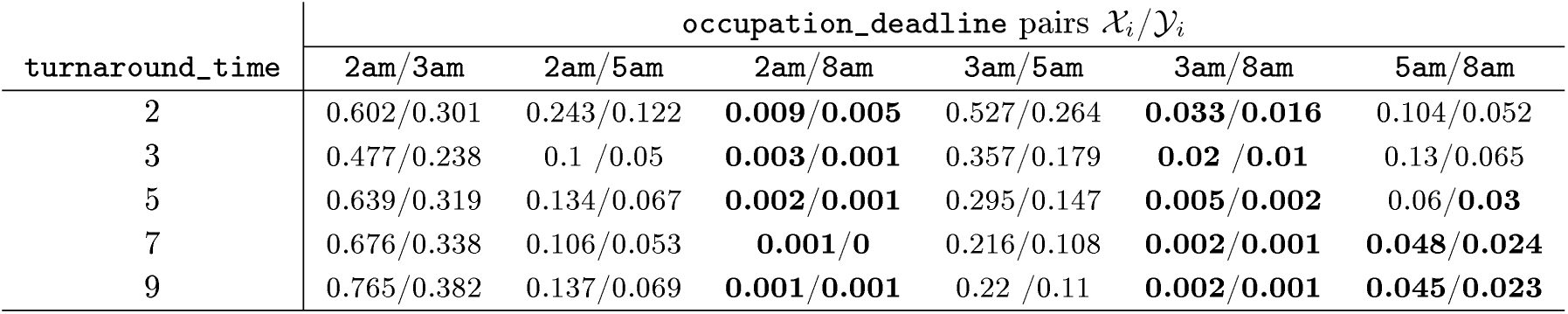
GB2 colony in 2008. Wilcox rank-sum test with 10^3^ bootstraps and `lf_delay`=3 minutes. Table cells are formatted as *p*_1_/*p*_2_ where *p*_1_ and *p*_2_ are the p-values for the hypotheses *ℋ*_1_ and *ℋ*_2_ respectively (see main text).

Here, `lf_delay` is fixed at 3 minutes, while `occupation_deadline` is varied in {2am, 3am, 5am, 8am}, and `turnaround_time` - in {2, 3, 5, 7, 9} minutes. For each value of turnaround_time (rows in the table), we compare the time difference distributions (*𝒳*_*i*_/*𝒴*_*i*_) between all possible pairs of `occupation_deadline`. The comparison is done via a bootstrapped Wilcoxon rank-sum test on the null hypothesis that the two distributions are the same, against the two-sided alternative *ℋ*_1_, and the one-sided alternative *ℋ*_2_ that *𝒳*_*i*_ < *𝒴*_*i*_. Each table cell shows the p-value for the two-sided and one-sided test, respectively.

As an example, fixing `turnaround_time` = 2 minutes, we see that the distribution of L/F time differences for `occupation_deadline` at 2am is not statistically different from the distribution with `occupation_deadline` at 3am (p-value = 0.602). This is an indication that the nature of the identified L/F events is invariant to the later deadline, hence it is unlikely that we have inadvertently included swarming effects. Further inspection of the table reveals that qualitative changes in L/F time differences occur when `occupation_deadline`=8am, but not for the other pair-wise comparisons. The one-sided test indicates the type of these changes, namely that L/F events inferred up to 8am on the day of occupation, tend to have larger time differences compared to earlier occupation deadlines. This is in line with the reasoning in Appendix S.4 and implies the presence of swarming effects. Therefore, `occupation_deadline`=8am is likely too late.

Moreover, this conclusion holds when varying `turnaround_time`, as well. The impact of this parameter on the L/F time differences seems to be small, in the range considered, except for values smaller than 5 minutes and comparing `occupation_deadline` = 5am vs. `occupation_deadline` =8am. In these cases, too many events with small time differences are identified, which conceals the swarming events. The effect of `turnaround_time` is primarily on the number of identified L/F events, as assuming larger recruitment delays excludes events where the leader found a follower relatively quickly (Table **S5**).

Based on these arguments, for a fixed `lf_delay`=3 minutes, we would choose `turnaround_time`=3 minutes and `occupation_deadline`=5am on the day of occupation.

This gives us an optimal trade-off between the number of inferred L/F events, and the interference due to swarming.

In Table **S4** we apply the same comparison procedure, but this time we fix `lf_delay` =5 minutes. Again, `occupation_deadline` =8am produced consistently larger time differences that are not present when comparing all other `occupation_deadline` pairs. Additionally, the effect of `turnaround_time` is again small. Considering that higher `lf_delay` further increases our sample of identified L/F events (Table S5), we fix `lf_delay` =5 minutes.

**Table S4:** GB 2 colony in 2008 with `lf_delay` =5 minutes.

| turnaround_time | occupation_deadline pairs |  |  |  |  |  |
| --- | --- | --- | --- | --- | --- | --- |
|  | 2am/3am | 2am/5am | 2am/8am | 3am/5am | 3am/8am | 5am/8am |
| 2 | 0.725/0.362 | 0.522/0.261 | **0.005/0.003** | 0.782/0.391 | **0.011/0.006** | **0.012/0.006** |
| 3 | 0.619/0.31 | 0.349/0.175 | **0.006/0.003** | 0.671/0.335 | **0.019/0.009** | **0.03/0.015** |
| 5 | 0.457/0.229 | 0.135/0.068 | 0/0 | 0.47/0.235 | **0.004/0.002** | **0.018/0.009** |
| 7 | 0.457/0.228 | 0.094/**0.047** | 0/0 | 0.36/0.18 | **0.002/0.001** | **0.015/0.008** |
| 9 | 0.514/0.257 | 0.085/**0.043** | 0/0 | 0.29/0.145 | **0.001/0** | **0.012/0.006** |

**Table S5:**
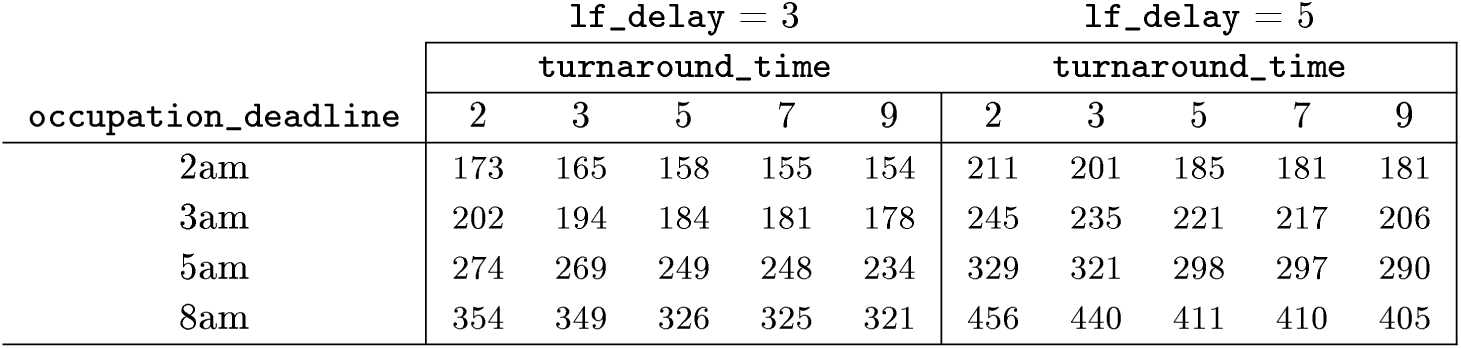
Number of identified L/F events for the GB2 colony in 2008 with different values of the three parameters.

### S.4 Effects of swarming on L/F time differences

Below we illustrate how swarming affects L/F time differences. In particular, the time differences, in presence of swarming, are skewed towards the **lf_delay** limit. Table S6 shows a typical recording pattern from a representative experimental box close to 6am on the day of the box occupation. Here, five experienced and five naïve individuals were recorded within about 10 minutes. Table **S7** contains all L/F events identified from the sample with **lf_delay** fixed at 5 minutes.

How do we identify L/F events from such data, without confounding them with swarming behavior? In fact, we are not interested in identifying swarming behavior. Instead, we only want to extract legitimate L/F events. In Section S.3, we already analyzed the distributions of time differences between L/F events, to detect for which parameter values the distributions start to deviate significantly. This is sufficient to calibrate the three parameters of our method.

**Table S6:**
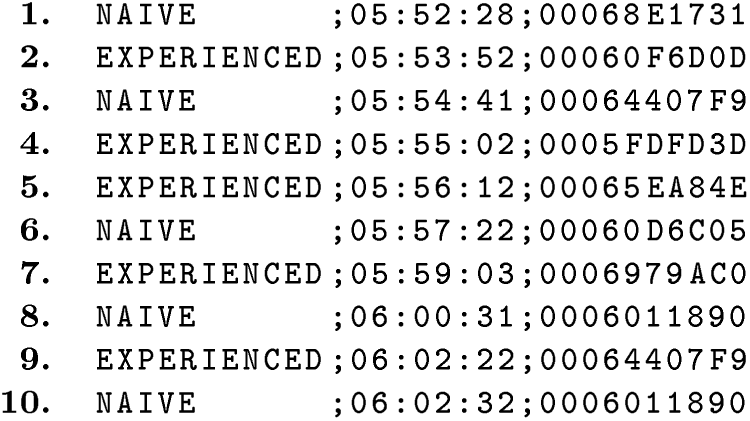
An excerpt from the *processed* recordings of an experimental box for the GB2 colony in 2008. Each line contains the individual information status, recording time, and unique identification number, in that order, separated by semi-colon.

Here we also offer a possible explanation for these deviations. In Table S6, the mean time difference of the events is 2.4 minutes, and the minimum is strictly above 1 minute. We argue that this characteristic is more consistent with swarming behaviour, in which a few experienced individuals attract naïve conspecifics by circling around the roost and emitting echolocation calls. Since experienced and naïve individuals are not grouped together as in genuine leading-following pairs, it takes time for a naïve individual to respond to the calls and fly to the roost. As a result, most L/F events identified in this way tend to have larger time differences closer to the allowable limit of **lf_delay**. We use precisely this observation when fine-tuning the **occupation_deadline** parameter.

As a comparison, consider the recording pattern in Table S9 from another box close to 2am on the day of its occupation. The L/F events corresponding to this pattern are shown in Table **S8**. The mean time difference is 1.5 minutes and the minimum is zero, as individual arrivals exceeded the time resolution of the reading device. This indicates that an experienced individual did appear close together with a designated follower. As for the couple of events with large time differences, they are most likely due to naïve individuals remaining at the entrance of the box, thereby triggering the reading device repetitively, than to swarming. As seen from Table S9, three such individuals, **00065DB1F6, 000697D00F** and **00068E1B66**, generate long recording sequences that prevent other followers from examining the box upon arrival, thereby forming an L/F event with large time difference to the leader.

**Table S7:** The L/F events corresponding to the recording pattern in Table S6. Parameters `lf_delay`=5 minutes and `occupation_deadline`=8am. `turnaround_time` does not affect this example, as no naïve individual appears again as a leader.

| Follower | Leader | L/F time<br>difference | L/F times |
| --- | --- | --- | --- |
| 00060F6D0D | 00068E1731 | 1.4 | 05:53:52 / 05:52:28 |
| 00064407F9 | 00068E1731 | 2.21667 | 05:54:41 / 05:52:28 |
| 0005FDFD3D | 00068E1731 | 2.56667 | 05:55:02 / 05:52:28 |
| 00065EA84E | 00068E1731 | 3.73333 | 05:56:12 / 05:52:28 |
| 00060F6D0D | 00060D6C05 | 3.5 | 05:53:52 / 05:57:22 |
| 00064407F9 | 00060D6C05 | 2.68333 | 05:54:41 / 05:57:22 |
| 0005FDFD3D | 00060D6C05 | 2.33333 | 05:55:02 / 05:57:22 |
| 00065EA84E | 00060D6C05 | 1.16667 | 05:56:12 / 05:57:22 |
| 00065EA84E | 0006011890 | 4.31667 | 05:56:12 / 06:00:31 |
| 0006979AC0 | 00060D6C05 | 1.68333 | 05:59:03 / 05:57:22 |
| 0006979AC0 | 0006011890 | 1.46667 | 05:59:03 / 06:00:31 |
| 00064407F9 | 0006011890 | 1.85 | 06:02:22 / 06:00:31 |

**Table S8:**
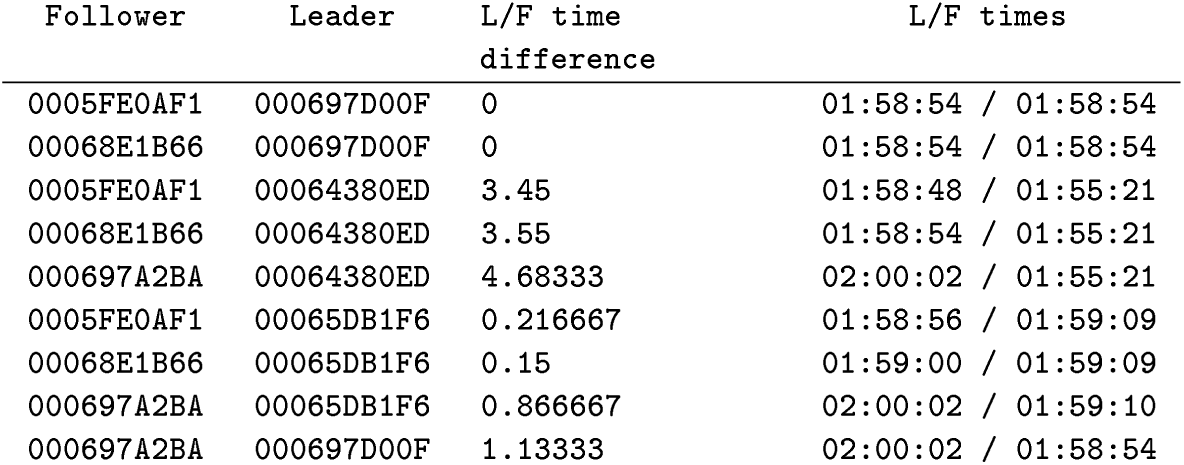
The L/F events corresponding to Table S9. Parameters as in Table **S7**.

**Table S9:** An excerpt from the *processed* recordings of an experimental box for the GB2 colony in 2008. The pattern is formatted as in Table S6.

|  |  |  |
| --- | --- | --- |
| 1. | NAIVE | ;01:55:21;00064380ED |
| 2. | NAIVE | ;01:55:25;00065DB1F6 |
| 3. | NAIVE | ;01:55:26;00065DB1F6 |
| 4. | NAIVE | ;01:55:36;000697D00F |
| 5. | NAIVE | ;01:55:49;00065DB1F6 |
| 6. | NAIVE | ;01:55:50;00065DB1F6 |
| 7. | NAIVE | ;01:55:51;00065DB1F6 |
| 8. | NAIVE | ;01:55:52;00065DB1F6 |
| 9. | NAIVE | ;01:55:55;00065DB1F6 |
| 10. | NAIVE | ;01:57:12;00065DB1F6 |
| 11. | NAIVE | ;01:57:14;00065DB1F6 |
| 12. | NAIVE | ;01:57:21;000697D00F |
| 13. | NAIVE | ;01:57:22;000697D00F |
| 14. | NAIVE | ;01:57:23;000697D00F |
| 15. | NAIVE | ;01:57:25;000697D00F |
| 16. | NAIVE | ;01:57:27;000697D00F |
| 17. | NAIVE | ;01:58:30;000697D00F |
| 18. | NAIVE | ;01:58:31;000697D00F |
| 19. | NAIVE | ;01:58:32;000697D00F |
| 20. | NAIVE | ;01:58:35;000697D00F |
| 21. | NAIVE | ;01:58:36;000697D00F |
| 22. | NAIVE | ;01:58:45;000697D00F |
| 23. | NAIVE | ;01:58:46;000697D00F |
| 24. | NAIVE | ;01:58:47;000697D00F |
| 25. | EXPERIENCED | ;01:58:48;0005FE0AF1 |
| 26. | EXPERIENCED | ;01:58:49;0005FE0AF1 |
| 27. | EXPERIENCED | ;01:58:50;0005FE0AF1 |
| 28. | EXPERIENCED | ;01:58:52;0005FE0AF1 |
| 29. | NAIVE | ;01:58:53;000697D00F |
| 30. | EXPERIENCED | ;01:58:54;0005FE0AF1 |
| 31. | EXPERIENCED | ;01:58:54;00068E1B66 |
| 32. | NAIVE | ;01:58:54;000697D00F |
| 33. | EXPERIENCED | ;01:58:55;0005FE0AF1 |
| 34. | EXPERIENCED | ;01:58:56;0005FE0AF1 |
| 35. | EXPERIENCED | ;01:58:56;00068E1B66 |
| 36. | EXPERIENCED | ;01:58:57;00068E1B66 |
| 37. | EXPERIENCED | ;01:58:58;00068E1B66 |
| 38. | EXPERIENCED | ;01:58:59;00068E1B66 |
| 39. | EXPERIENCED | ;01:59:00;00068E1B66 |
| 40. | NAIVE | ;01:59:09;00065DB1F6 |
| 41. | NAIVE | ;01:59:10;00065DB1F6 |
| 42. | EXPERIENCED | ;02:00:02;000697A2BA |

### S.5 Eigenvector centrality explained

Consider the following simple leading-following network with 4 bats (nodes) and 5 leading-following events (links):

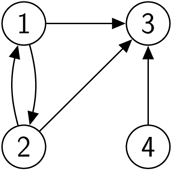

An alternative representation of this network is through its so called *adjacency* matrix, *A*, which indicates which two nodes are adjacent to each other.

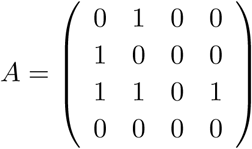

The elements *a*_*i,j*_ (*i*, and *j* index rows and columns, respectively) in this matrix are 1 if a directed link exists between nodes *j* and *i*. In other words, *a*_*i,j*_ = 1 if *j* followed *i*. Otherwise, *a*_*i,j*_ = 0. For example the first column *a*_*i*,1_ gives all nodes that node 1 follows. We see that *a*_2,1_ = *a*_3,1_ = 1, so 1 has followed both 2 and 3.

The main idea behind eigenvector centrality is that the centrality of a node *i, c*_*i*_, is proportionate to the sum of the centralities of all nodes who follow it. Staying with node 1, its centrality is the sum of the centralities of nodes 2 and 3, i.e. 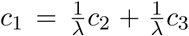 or *λ.c*_1_ = *c*_2_ + *c*_3_ for some proportionality constant *λ*. In this way, we can express the centralities of all nodes and write them as a system of equations:

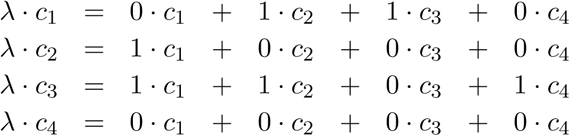

In matrix form the above system can be rewritten as:

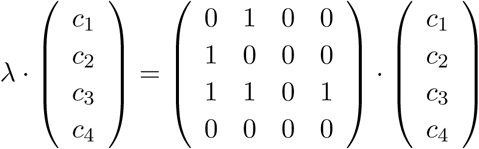

or in vector notation: 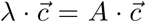. This is the familiar eigenvector problem. We need to find a vector 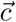 such that upon applying matrix *A* to it, the result is a scaled version of 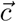 with a scaling factor *λ*. The unknown vector 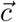 is called an eigenvector of the matrix *A*, and *λ* is referred to as the eigenvalue, which corresponds to that eigenvector. Solving the system of equations yields: 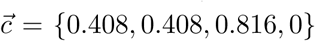 and *λ* = 1. Therefore node 3 is most central since it is followed by everyone. Nodes 1 and 2 follow each other so they boost their own centrality, and node 4 is not followed by anyone so its centrality is 0.

### S.6 Calculation of individual centrality measures

**Figure S2:**
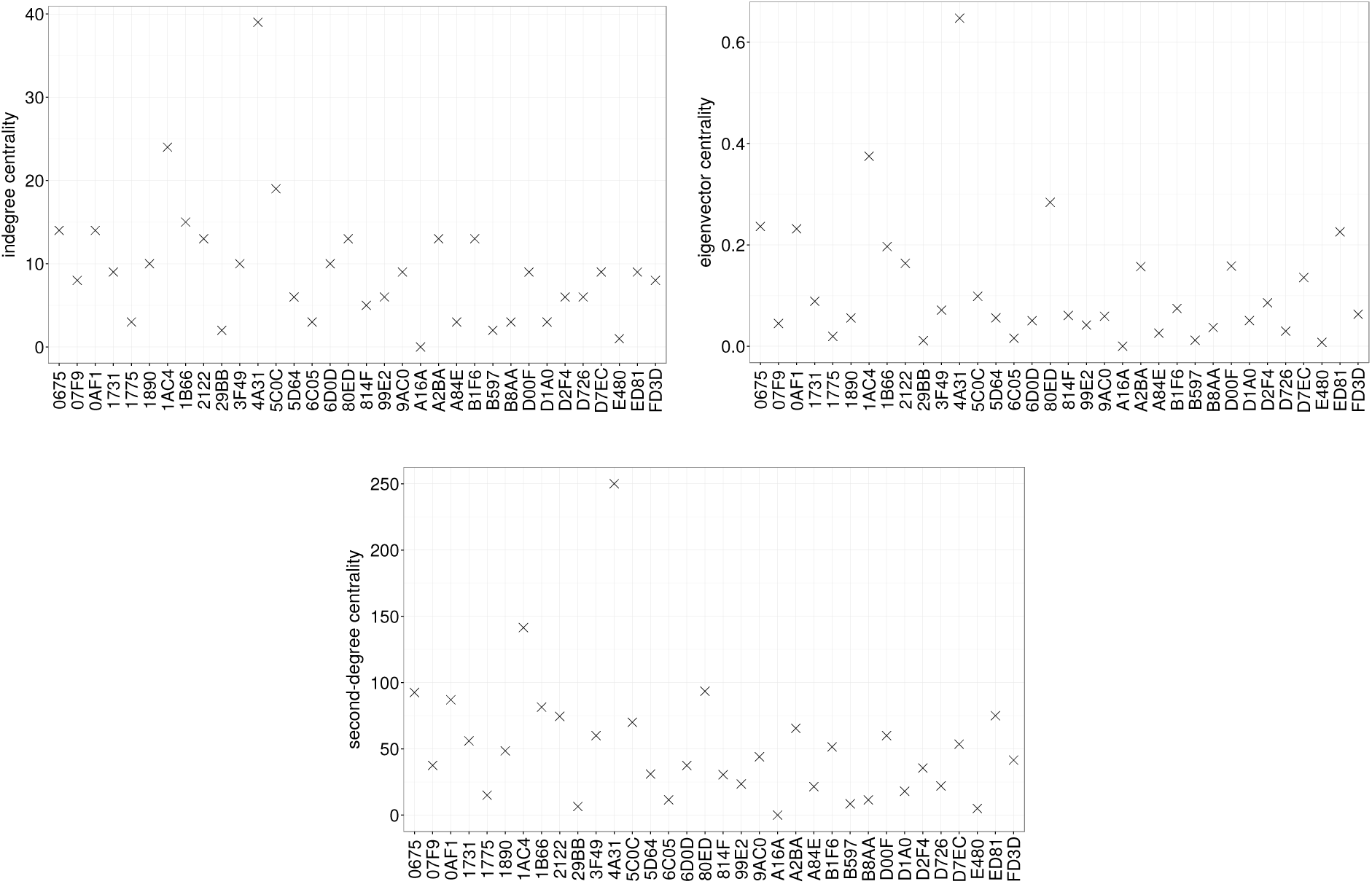
Individual influence quantified according to the three centrality measures introduced in Section 3.2: (top left) in-degree centrality, (top right) eigenvector centrality, (bottom) second-degree centrality (top left). For the calculation, the L/F network constructed from the dataset of colony GB2 in year 2008 was used. The x-axis displays the last four digits of a bat’s unique identification number.

### S.7 Correlations between centrality measures

**Table S10:** Correlations between the rankings produced by the three different centrality measures. *S* stands ranking using second degree centrality, *E* stands for ranking using eigenvector centrality, and *D* stands for ranking using in-degree centrality. The first column lists the different datasets as described in Table 2.

| | $S \Leftrightarrow E$ | $S \Leftrightarrow D$ | $D \Leftrightarrow E$ |
| --- | --- | --- | --- |
| GB2 2008 | 0.98 | 0.97 | 0.90 |
| GB2 2011 | 0.99 | 0.93 | 0.92 |
| BS 2007 | 0.99 | 0.98 | 0.98 |
| BS 2009 | 0.99 | 0.97 | 0.98 |
| BS 2010 | 0.99 | 0.96 | 0.94 |
| BS 2011 | 0.99 | 0.93 | 0.94 |

## Footnotes

† Objective, as in best reflection of reality. Indeed, one cannot “ask” a bat how much time she needs for recruitment or how far away she travels from a follower.

